# Altered structural hub connectivity and its clinical relevance in glioma

**DOI:** 10.1101/610618

**Authors:** Linda Douw, Julie J. Miller, Martijn D. Steenwijk, Steven M. Stufflebeam, Elizabeth R. Gerstner

## Abstract

**Background and Purpose:** Structural network analysis of diffusion imaging is increasingly used to study neurological disease, its pathophysiology and symptoms. We therefore evaluate structural hub connectivity in glioma patients and its association with molecular subtype and clinical status.

**Materials and Methods:** Using retrospective diffusion imaging, structural connectivity was investigated in 65 newly diagnosed glioma patients (36 males; mean age 52 ± 14 years) and 60 healthy controls (23 males; mean age 50 ± 7 years). Probabilistic tractography was performed between 39 cortical nodes per hemisphere. In patients, tumors were drawn in to exclude each tumor-containing voxel from analysis. As previous connectomic research in glioma and other neurological diseases has shown particular importance of ‘hub’ nodes and connections, the numbers of connections between hubs, hubs and non-hubs, and non-hubs were calculated for each hemisphere separately. Clinical and molecular characteristics were assessed as part of routine clinical care. Group differences in connectivity and its associations with performance and molecular subtypes were tested non-parametrically through Mann-Whitney U-tests, corrected for multiple comparisons.

**Results:** Glioma patients had more hub-related connections in the hemisphere contralateral to the tumor (hub-hub *P* = 0.002, hub-non-hub *P* = 0.005), despite being comparable to controls in terms of total and ipsilateral connections. Within patients, hub-related connectivity related to performance status (*P* = 0.009) and molecular subtype (*P* = 0.045).

**Conclusion:** We present experimental evidence for the relevance of structural connectomics as a tool to pick up on the clinical impact of glioma on the rest of the brain.

## Introduction

The neurological impact of pathology is increasingly understood in terms of dysfunctional groups of interconnected regions, as per the framework of network theory.^1,2^ In glioma, widespread functional connections are altered, correlating with patient functioning.^3–8^ Network ‘hubs’, i.e. extensively connected regions, seem most vulnerable to this dysfunction.^4,6–9^

Network theory also offers mathematical insight on how particularly increased hub-related connectivity relates to later large-scale network failure,^10,11^ and may therefore explain clinical decline that cannot be explained by currently used radiological features across several neurological diseases.^2,12–14^ In newly diagnosed glioma, hub-related functional connectivity in the hemisphere contralateral to the tumor is increased,^7^ possibly reflecting network failure in response to local dysfunction as an initially compensatory but long-term detrimental relaying of load.^10^ Structural connectivity throughout macroscopically tumor-free brain regions also experimentally associates with molecular subtype^15^ and prognosis,^16^ indicating that connectivity measures may inform our understanding of performance status and survival beyond the currently known molecular determinants.

We retrospectively investigated structural connectivity in newly diagnosed glioma patients, assessing altered structural connectivity, its associations with functioning and survival, and differences between molecular subtypes. We hypothesized that higher hub-non-hub-related connectivity would reflect lower performance status, shorter survival, and unfavorable molecular subtype, while maintained hub-connectivity would associate with better performance, longer survival, and favorable molecular subtype.

## Materials and Methods

### Participants

Newly diagnosed glioma patients that underwent diffusion imaging (dMRI) and T1-3D imaging at the time of diagnosis between January 2006 and December 2013 were retrospectively included. Inclusion criteria were histologically confirmed glioma, dMRI available before diagnostic surgery and/or treatment, and T1-3D imaging at the same time point. Exclusion criteria were previous craniotomy, previous chemo/radiotherapy, bilateral tumor invasion or significant midline shift, and neurological or psychiatric comorbidity, although presence of seizures was allowed given the frequency of seizures in glioma patients. The Institutional Review Board approved the retrospective analysis of patient data used, and we adhered to HIPAA regulations dealing with these data. As a healthy control group, we used a cohort that was collected in a different hospital. Prospective inclusion was approved by the Institutional Review Board; all controls gave written informed consent before participation.

Clinical information was extracted by medical chart review. Karnofsky Performance Status (KPS) was summarized categorically as 90-100 or <80.^7,15^ Progression-free survival (PFS) reflected the number of weeks between date of diagnosis and date of radiological/clinical progression. Overall survival (OS) reflected the number of weeks between date of diagnosis and date of death. Patients who had not progressed or died at final analysis (July 2016) were censored as of the last contact date with their treating neuro-oncologist.

### Molecular subtypes

The revised glioma classification of the World Health Organization establishes several relevant molecular markers,^17^ including isocitrate dehydrogenase (IDH1) mutations, tumor protein 53 (*TP53*) mutations, O6-methylguanin-DNA-methyltransferase (MGMT) methylation status, and 1p/19q codeletion. More favorable prognoses are related to presence of IDH1 mutation, 1p/19q codeletion, absence of *TP53* mutation, and MGMT methylation.^18–22^ Molecular subtypes were determined as part of routine clinical care and varied in availability (see results section).

### MRI

The diffusion sequence used in patients consisted of 21 diffusion-encoding gradient directions, and a b-value of 1000 (TR=5400ms, 1.38×1.38mm^2^ in-plane resolution). Healthy controls underwent imaging with 30 gradient directions at a b-value of 900 (TR=13000ms, 2×2mm^2^ in-plane resolution).

Figure 1 schematically depicts our analysis pipeline. Imaging processing steps were performed using FSL 5.0.9 (http://www.fmrib.ox.ac.uk/fsl). Non-brain tissue was removed using the Brain Extraction Tool,^23^ tissue segmentation was performed using FAST.^24^ The Automated Structural Labeling (AAL) atlas was used to define 78 cortical regions in each subject’s native space after registration with FLIRT.^25^ Tumor masks were created manually using contrast enhancing T1-weighted and FLAIR images. Tumor volume was calculated using this mask.

**Figure 1.**
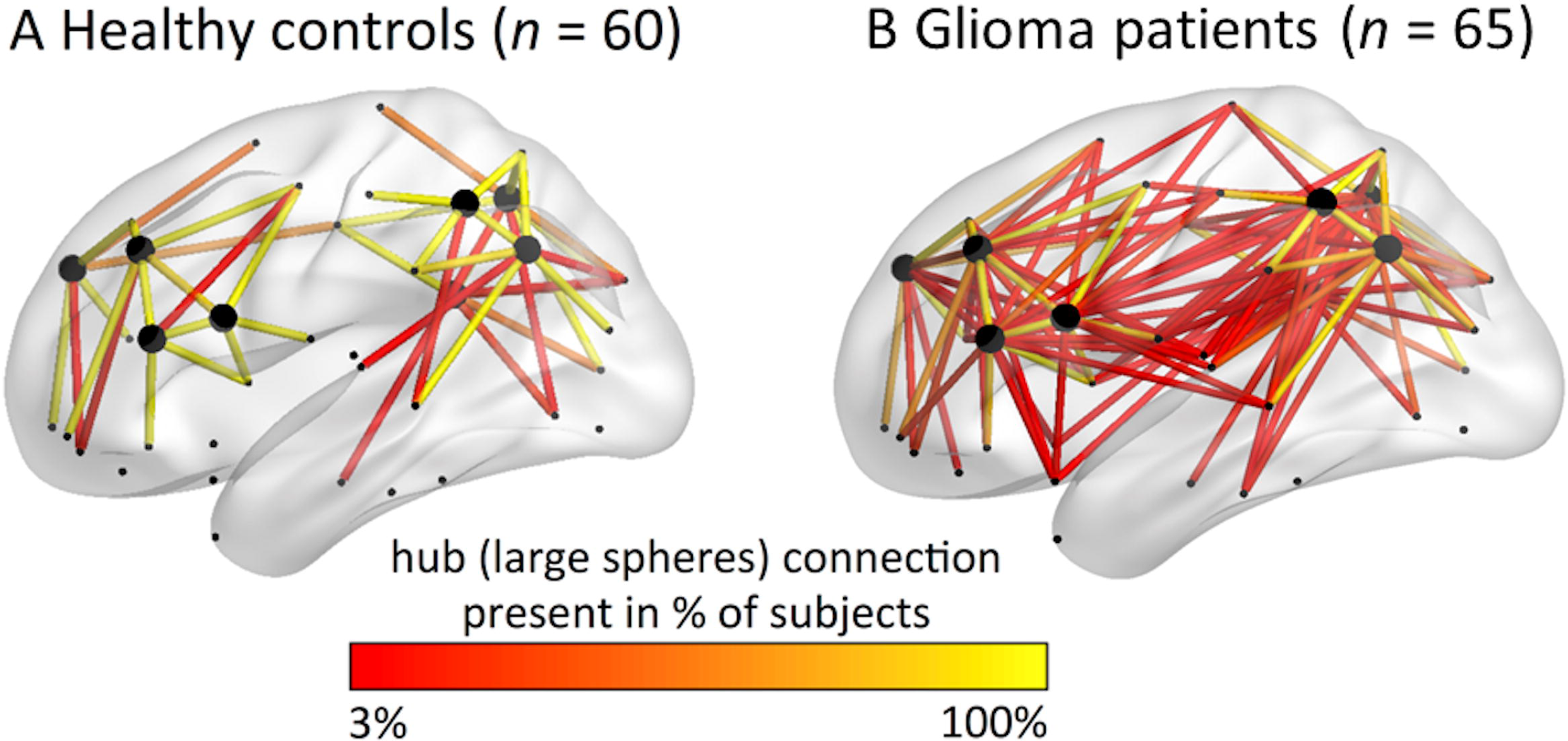
The analysis pipeline. In panel (A), an exemplar patient MRI is shown on the left according to neurological convention, with the tumor drawn in in the middle column, and the 78 region atlas spanning the cortical ribbon projected onto the brain on the right. In (B), we show an exemplar diffusion image on the left, such an image with the atlas regions spanning the grey/white matter rim in the middle, and the results of probabilistic tractography of the right primary motor cortex to the rest of the brain on the right. Panel (C) indicates hub regions and connections. On the left, the median connectome of the healthy controls is shown, with hub regions indicated in pink and the rest (non-hub regions) in blue. On the right of the panel, connections between hubs (large spheres) are indicated in pink, while connections between hubs and non-hubs (small spheres) are displayed in blue. Connections between non-hubs are not displayed, for clarity.

Diffusion images were visually inspected for artifacts and/or excessive motion, then corrected for motion and eddy current distortion using FMRIB’s Diffusion Toolbox,^26^ which was also used to fit the voxelwise diffusion tensor. Diffusion images were co-registered to individual T1s with epi_reg. Voxels forming the grey/white matter rim were seeds in subsequent probabilistic tractography (ProbtrackX2, 5000 streamlines per voxel). Tract likelihood was determined by averaging the number of streamlines reaching each atlas region from each other region across all voxels (excluding all tumor mask voxels), then normalizing this value for seed region size.

### Hub-related connectivity

Connectivity analyses were performed using Matlab R2012a (Mathworks, Natick, MA, US). Binary connectivity matrices were created by thresholding the number of normalized streamlines reaching their target in >5% of runs. In controls, the median density (i.e. number of connections) of the patients was used to threshold the weighted average of the left and right hemisphere control matrices, in order to keep the total number of connections comparable between patients and controls.

Hub locations were in accordance with the literature, encompassing eight regions falling within the default mode^27,28^ and frontoparietal networks.^29,30^ This ensured that spatial shifts in hub regions in patients would not drive our findings concerning alterations in connectivity of the well-known hubs of the human brain. Finally, we calculated the total number of connections per hemisphere, as well as the number of connections between hubs, hubs and non-hubs, and non-hubs for each subject.

### Statistical analysis

Matlab and SPSS 22.0 (IBM Corp, Armonk, NY, US) were used with an alpha level of 0.05. Patient-control characteristics were tested using Chi-square tests (sex) and Student’s t-test for independent samples (age).

Connectivity was non-normally distributed according to Kolmogorov-Smirnov tests. Differences in the number of connections between patients and controls were explored using Wilcoxon signed rank tests, corrected for eight comparisons using Bonferroni correction (significance level *P* < 0.006). Associations between significantly altered connectivity and KPS were tested using Mann-Whitney U-tests, while associations between hub connectivity and survival were investigated using log-rank tests, both with Bonferroni correction.

Differences in hub-related connectivity depending on molecular subtype were explored using Mann-Whitney U-tests. An uncorrected *P* < 0.05 was considered statistically significant for these exploratory analyses.

## Results

### Subject characteristics

In total, 293 newly diagnosed, preoperative glioma patients were screened for inclusion. One patient was excluded because of neurological comorbidities, 69 and 82 due to absence of dMRI or 3D-T1 imaging, respectively, 38 due to scanning artifacts, while another 39 patients underwent dMRI with different scanning sequences, leaving 65 patients for final analysis. We found no significant differences in age or sex between patients and controls (Table 2).

**Table 1.** Patient characteristics

| Variable | Patients (*n* = 65) |
| --- | --- |
| Mean age in years (SD) | 52 (14) |
| Males (females) | 36 (29) |
| KPS 90-100 \| KPS < 90 \| NA | 42 \| 18 \| 5 |
| WHO 2007 glioma grade II \| III \| IV | 8 \| 4 \| 53 |
| Left (right) tumor lateralization | 34 (31) |
| AED users \| non-users \| NA | 46 \| 17 \| 2 |
| EOR gross total \| subtotal \| biopsy \| NA | 29 \| 25 \| 10 \| 1 |
| Median tumor volume in cm<sup>3</sup> (IQR) | 26.7 (49.0) |
| 1p/19q codeletion \| no codeletion \| NA | 2 \| 27 \| 36 |
| IDH mutation (WT) | 7 (3) |
| TP53 mutation (WT) | 7 (58) |
| MGMT methylated \| unmethylated \| NA | 26 \| 22 \| 17 |

Note. KPS = Karnofsky Performance Status, WHO=World Health Organization, AED = antiepileptic drugs, EOR = extent of resection, IDH = isocitrate dehydrogenase, TP53 = tumor protein 53, MGMT = O6-methylguanin-DNA-methyltransferase, NA = not available.

**Table 2.** Characteristics and connectivity in patients versus controls

| Variable | Controls (*n* = 60) | Patients (*n* = 65) | Test statistic | *P* value |
| --- | --- | --- | --- | --- |
| Mean age in years (SD) | 50 (7) | 52 (14) | $t(123) = 1.10$ | 0.31 |
| Number of males (females) | 23 (37) | 29 (36) | $\chi^2 = 3.64$ | 0.06 |
| Mdn ipsi (IQR) | 194 (0) | 194 (55) | $U = 1950$ | 0.99 |
| Mdn ipsi hub (IQR) | 10 (0) | 10 (2) | $U = 1725$ | 0.16 |
| Mdn ipsi hub-non-hub (IQR) | 56 (4) | 60 (20) | $U = 1555$ | 0.01 |
| Mdn ipsi non-hub (IQR) | 128 (4) | 120 (39) | $U = 2176$ | 0.26 |
| Mdn contra (IQR) | 194 (0) | 194 (63) | $U = 2010$ | 0.75 |
| Mdn contra hub (IQR) | 10 (0) | 10 (3) | $U = 1460$ | 0.002* |
| Mdn contra hub-non-hub (IQR) | 56 (4) | 60 (19) | $U = 1387$ | 0.005* |
| Mdn contra non-hub (IQR) | 128 (4) | 122 (38) | $U = 2216$ | 0.19 |

Note. \* = $P < 0.05$ after correction for multiple comparisons, Mdn = median, ipsi = ipsilateral to the tumor, contra = contralateral to the tumor, IQR = interquartile range.

### Patients had more contralateral hub-related connections than controls

There were no significant differences between patients and healthy controls in ipsilateral or contralateral connection density, nor did patients have an altered number of non-hub connections in either hemisphere (Table 2 and electronic supplementary material). Furthermore, the number of hub-related connections in the ipsilateral hemisphere was not significantly different between groups. However, patients did have a higher number of connections between hubs (U = 1460, *P* = 0.002), and between hubs and non-hubs (U = 1387, *P* = 0.005) in the contralateral hemisphere (Figure 2). Within patients, we explored a number of possible confounders of these contralateral hub indices. The number of contralateral hub-hub connections did not relate to age (Tau = 0.15, *P* = 0.10), tumor volume (Tau = 0.09, *P* = 0.35), the extent of overlap between the tumor and the hubs (Tau = 0.02, *P* = 0.86), tumor grade according to the 2007 WHO grading system (Kruskal-Wallis = 2.51, *P* = 0.29), AED use (Mann-Whitney U = 464, *P* = 0.22), or corticosteroid use (Mann-Whitney U = 460, *P* = 0.56). The number of contralateral hub-non-hub connections relate to age (Tau = 0.09, *P* = 0.30), tumor volume (Tau = 0.04, *P* = 0.67), hub overlap (Tau = 0.07, *P* = 0.45), tumor grade (Kruskal-Wallis = 1.02, *P* = 0.60), AED use (Mann-Whitney U = 454, *P* = 0.33), or corticosteroid use (Mann-Whitney U = 480, *P* = 0.79).

**Figure 2.**
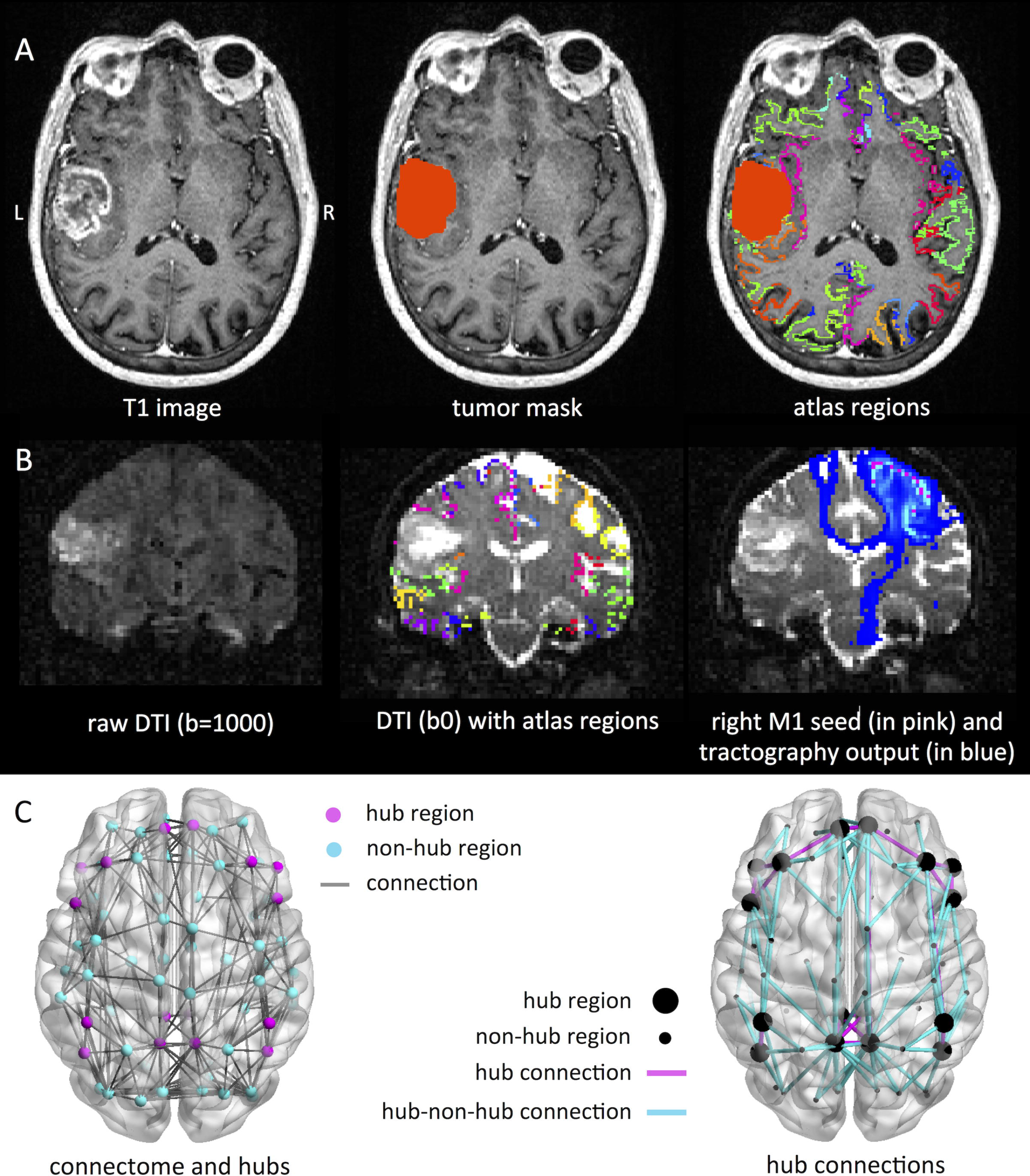
Hub-related connectivity in patients versus controls. All hub-related connections are depicted on this lateral view of the brain, for (A) patients and (B) healthy controls. Hub nodes are depicted as larger spheres than non-hub nodes. The color of each connection indicates the percentage of subjects in which this connection was present within each group.

### Higher hub-non-hub connectivity was associated with poorer performance status but not survival

Patients with low KPS had higher contralateral hub-non-hub connectivity (median 75 connections, IQR 39) than patients with favorable KPS (median 58 connections, IQR 29; U = 216, *P* = 0.009). The number of contralateral hub connections was not significantly related to KPS (U = 259, *P* = 0.04). Hub-non-hub connectivity did not relate to either PFS (Chi-square = 0.05, *P* = 0.82) or OS (Chi-square = 0.004, *P* = 0.95), nor did the number of contralateral hub connections (PFS Chi-square = 0.77, *P* = 0.378; OS Chi-square = 0.18, *P* = 0.67).

### Higher hub-hub connectivity related to favorable molecular subtype

Without correction for multiple comparisons, non-GBM IDH mut patients had a significantly higher number of contralateral hub connections than IDH wt patients (see table 3). There were no differences in hub-non-hub connections according to molecular subtypes.

**Table 3.** Connectivity profiles within molecular subtypes

| Molecular subtypes | Mdn contra<br>hub, IQR | Test<br>statistic | *P* value | Mdn contra<br>hub-non-hub,<br>IQR | Test<br>statistic | *P*<br>value |
| --- | --- | --- | --- | --- | --- | --- |
| Non-GBM |  |  |  |  |  |  |
| IDH mut, *n* = 6 (wt, *n* = 3) | 10, 1 (8, 1) | U = 16 | 0.045* | 55, 24 (62, 18) | U = 9 | 0.99 |
| TP53 mut, *n* = 3 (wt, *n* = 9) | 10, 0 (10, 3) | U = 14 | 0.99 | 52, 4 (62, 18) | U = 8 | 0.31 |
| GBM |  |  |  |  |  |  |
| IDH mut, *n* = 0 (wt, *n* = 0) | NA | NA | NA | NA | NA | NA |
| TP53 mut, *n* = 4 (wt, *n* = 49) | 15, 8 (10, 2) | U = 150 | 0.06 | 67, 30 (60, 22) | U = 111 | 0.661 |

Note. \* $P < 0.05$ (no correction for multiple comparisons). NA = not applicable due to small sample
sizes, Mdn = median, contra = contralateral to the tumor, IQR = interquartile range, U = Mann-
Whitney test statistic, TP53 = tumor protein 53, IDH = isocitrate dehydrogenase.

## Discussion

We report higher hub-related structural connectivity in glioma patients in the hemisphere contralateral to the tumor, as compared to healthy controls. The total number of connections and number of (hub-related) connections in the macroscopically invaded hemisphere did not differ between groups, nor could altered hub-related connectivity be attributed to tumor grade, volume, overlap with hub regions, or AED and/or corticosteroid use. Furthermore, higher contralateral hub-non-hub connectivity related to poorer performance status, while higher contralateral hub-hub connectivity related to a favorable molecular subtype in terms of IDH mutation status.

Increased hub-related connectivity may speculatively support the hypothesis that these patients are in the first phase of cascadic network failure, as has been proposed and experimentally investigated in other neurological diseases.^2,12^ Connectivity between non-hubs and hubs is thought to increase due to local dysfunction and subsequent relaying of connectivity towards the network backbone,^10,11,13^ as confirmed by increased hub-related functional connectivity in newly diagnosed glioma patients.^7^ Conversely, maintained (or in this case: higher) hub-hub connectivity may indicate that the entire connectome has not collapsed yet, which would indicate the chronic phase of hub overload.

Previous studies have reported on (long-distance) reductions in connectivity, without exploring possible increases in structural connectivity.^31,32^ However, white matter integrity may increase at the scale of days in the setting of external brain stimulation.^33^ Our current results suggest that glioma is also able to induce higher white matter integrity at a distance from the tumor, be it a pathological sign of disease stage, compensatory (as suggested by the hub-hub association with favorable molecular subtype), or both. Of course, we cannot fully exclude the possibility that patients already had higher numbers of structural connections than controls before developing glioma, or that differing scanning protocols induces group differences between patients and controls. However, the within-patient associations found for both hub-non-hub connectivity (with unfavorable performance status) and hub-hub connectivity (with favorable mutation status) suggest that these factors do not confound the clinical implications of our findings. Futhermore, our results regarding lowered hub-hub connectivity in IDH wt patients corroborate a previous study of global structural network efficiency,^15^ as Kesler and colleagues show that IDH wt patients have reduced global efficiency as compared to IDH mut patients.

In the context of cascadic network failure, most previous work was performed using functional connectivity.^7,12^ Multimodal investigation of connectivity may prove essential to fully understanding the course of disease and performance status as a result of particular molecular subtypes of glioma. Also, it remains unclear how neuronal biology leads to circuit activity at the scale of thousands of cells, or how this circuitry precisely relates to macroscopic patterns of connectivity measured with MRI. We here chose to approach glioma from the complex network perspective in a top-down manner: if we assume that the macroscopic structural brain network adheres to mechanisms defined in other complex networks, and these mechanisms relate to relevant patient characteristics such as performance status and survival, future studies may use these findings to further uncover the underlying mechanisms from the bottom-up.

Another limitation of this study is its retrospective nature, limiting the availability of clinical information. Replication of our results in a prospective cohort of patients and controls is necessary. Our cross-sectional study also does not elucidate the timeframe in which connectivity alterations may have developed in glioma patients. Secondly, the boundaries of gliomas are challenging to define, so we cannot fully exclude carry-over tumor effects on contralateral diffusion measurements.

Furthermore, we used mostly functionally determined hub locations, instead of defining hubs based on (individual) structural data. Although the correlation between structural and functional connectivity has been shown particularly strong for the hub connections,^34^ future work may address the question whether the structural hubs we defined here are exactly identical to functional hubs on an individual level. Finally, optimal tractography methodology is currently debated upon, particularly due to false positives.^35^ However, we were primarily interested in hub connections, which are likely less influenced by weighted/proportional thresholding than non-hub connections, due to their overall higher strength and replicability. Furthermore, the use of a rather coarse atlas and limited number of diffusion gradients does imply greater clinical potential across a range of scanners.

## Conclusions

Our results reveal higher hub-related connectivity in glioma patients as compared to healthy controls, which may carry clinical information in terms of performance status and molecular subtype. We speculate that the co-occurrence of higher hub-related connectivity and lower performance status may indicate that these patients are in the first phase of hub overload. Conversely, maintained and/or higher hub-related connectivity may offer resilience in patients with favorable molecular subtypes. Our results offer a first step towards new avenues for the adequate understanding and possible future prognostic use of network failure in glioma patients.

## References

1. Crossley NA, Mechelli A, Scott J, et al. The hubs of the human connectome are generally implicated in the anatomy of brain disorders. Brain 2014;137:2382–95.

2. Aerts H, Fias W, Caeyenberghs K, et al. Brain networks under attack: robustness properties and the impact of lesions. Brain 2016;139:3063–83.

3. Bosma I, Reijneveld JC, Klein M, et al. Disturbed functional brain networks and neurocognitive function in low-grade glioma patients: a graph theoretical analysis of resting-state MEG. Nonlinear Biomed Phys 2009;3:9.

4. Esposito R, Mattei PA, Briganti C, et al. Modifications of default-mode network connectivity in patients with cerebral glioma. PLoS One 2012;7:e40231.

5. van Dellen E, Douw L, Hillebrand A, et al. MEG Network Differences between Low- and High-Grade Glioma Related to Epilepsy and Cognition. PLoS One 2012;7:e50122.

6. Harris RJ, Bookheimer SY, Cloughesy TF, et al. Altered functional connectivity of the default mode network in diffuse gliomas measured with pseudo-resting state fMRI. J Neurooncol 2014;116:373–9.

7. Derks J, Dirkson AR, de Witt Hamer PC, et al. Connectomic profile and clinical phenotype in newly diagnosed glioma patients. NeuroImage Clin https://doi.org/10.1016/j.nicl.2017.01.007.

8. Ghumman S, Fortin D, Noel-Lamy M, et al. Exploratory study of the effect of brain tumors on the default mode network. J Neurooncol 2016;128:437–44.

9. van Dellen E, Hillebrand A, Douw L, et al. Local polymorphic delta activity in cortical lesions causes global decreases in functional connectivity. Neuroimage 2013;83:524–32.

10. Buldyrev S V, Parshani R, Paul G, et al. Catastrophic cascade of failures in interdependent networks. Nature 2010;464:1025–8.

11. Motter AE, Yang Y. The unfolding and control of network cascades. Phys Today 2017;70:32–9.

12. Jones DT, Knopman DS, Gunter JL, et al. Cascading network failure across the Alzheimer’s disease spectrum. Brain 2016;139:547–62.

13. Stam CJ. Modern network science of neurological disorders. Nat Rev Neurosci 2014;15:683–95.

14. Jacobs HIL, Hedden T, Schultz AP, et al. Structural tract alterations predict downstream tau accumulation in amyloid-positive older individuals. Nat Neurosci 2018;21:424–31.

15. Kesler SR, Noll K, Cahill DP, et al. The effect of IDH1 mutation on the structural connectome in malignant astrocytoma. J Neurooncol https://doi.org/10.1007/s11060-016-2328-1.

16. Liu L, Zhang H, Rekik I, et al. Outcome Prediction for Patient with High-Grade Gliomas from Brain Functional and Structural Networks. In: Medical image computing and computer-assisted intervention◻: MICCAI … International Conference on Medical Image Computing and Computer-Assisted Intervention. Vol 9901.; 2016:26–34.

17. Louis DN, Perry A, Reifenberger G, et al. The 2016 World Health Organization Classification of Tumors of the Central Nervous System: a summary. Acta Neuropathol 2016;131:803–20.

18. Schiff D, Brown PD, Giannini C. Outcome in adult low-grade glioma: the impact of prognostic factors and treatment. Neurology 2007;69:1366–73.

19. Weller M, Stupp R, Hegi ME, et al. Personalized care in neuro-oncology coming of age: why we need MGMT and 1p/19q testing for malignant glioma patients in clinical practice. Neuro Oncol 2012;14:iv100–iv108.

20. Yan H, Parsons DW, Jin G, et al. IDH1 and IDH2 mutations in gliomas. N Engl J Med 2009;360:765–73.

21. Van Meir EG, Kikuchi T, Tada M, et al. Analysis of the p53 gene and its expression in human glioblastoma cells. Cancer Res 1994;54:649–52.

22. van den Bent MJ, Gravendeel LA, Gorlia T, et al. A Hypermethylated Phenotype Is a Better Predictor of Survival than MGMT Methylation in Anaplastic Oligodendroglial Brain Tumors: A Report from EORTC Study 26951. Clin Cancer Res 2011;17:7148–55.

23. Smith SM. Fast robust automated brain extraction. Hum Brain Mapp 2002;17:143–55.

24. Zhang Y, Brady M, Smith S. Segmentation of brain MR images through a hidden Markov random field model and the expectation-maximization algorithm. IEEE Trans Med Imaging 2001;20:45–57.

25. Tzourio-Mazoyer N, Landeau B, Papathanassiou D, et al. Automated anatomical labeling of activations in SPM using a macroscopic anatomical parcellation of the MNI MRI single-subject brain. Neuroimage 2002;15:273–89.

26. Smith SM, Jenkinson M, Woolrich MW, et al. Advances in functional and structural MR image analysis and implementation as FSL. Neuroimage 2004;23:S208–19.

27. Buckner RL, Sepulcre J, Talukdar T, et al. Cortical hubs revealed by intrinsic functional connectivity: mapping, assessment of stability, and relation to Alzheimer’s disease. J Neurosci 2009;29:1860–73.

28. van den Heuvel MP, Hulshoff Pol HE. Exploring the brain network: a review on resting-state fMRI functional connectivity. Eur Neuropsychopharmacol 2010;20:519–34.

29. Power JD, Schlaggar BL, Lessov-Schlaggar CN, et al. Evidence for hubs in human functional brain networks. Neuron 2013;79:798–813.

30. Cole MW, Reynolds JR, Power JD, et al. Multi-task connectivity reveals flexible hubs for adaptive task control. Nat Neurosci 2013;16:1348–55.

31. Mandonnet E, Cerliani L, Siuda-Krzywicka K, et al. A network-level approach of cognitive flexibility impairment after surgery of a right temporo-parietal glioma. Neurochirurgie 2017;63:308–13.

32. Bucci M, Mandelli ML, Berman JI, et al. Quantifying diffusion MRI tractography of the corticospinal tract in brain tumors with deterministic and probabilistic methods. NeuroImage Clin 2013;3:361–8.

33. Lindenberg R, Nachtigall L, Meinzer M, et al. Differential effects of dual and unihemispheric motor cortex stimulation in older adults. J Neurosci 2013;33:9176–83.

34. van den Heuvel MP, Mandl RC, Kahn RS, et al. Functionally linked resting-state networks reflect the underlying structural connectivity architecture of the human brain. Hum Brain Mapp 2009;30:3127–41.

35. Maier-Hein KH, Neher PF, Houde J-C, et al. The challenge of mapping the human connectome based on diffusion tractography. Nat Commun 2017;8:1349.

